# Detection of antibodies specific to H5 avian influenza virus in a sheep in Norway, June 2024, eleven months after an outbreak of highly pathogenic avian influenza in a nearby seabird colony

**DOI:** 10.1101/2025.08.14.670265

**Authors:** Johanna Hol Fosse, Grim Rømo, Francesco Bonfante, Ida Kristin Myhrvold, Kristin Stangeland Soetart, Kristin Udjus, Ragnhild Tønnessen

**Affiliations:** Norwegian Veterinary Institute, 1433 Ås, Norway; Istituto Zooprofilattico Sperimentale delle Venezie, Legnaro, Italy

**Author notes:** These authors contributed equally to this work and share first authorship (Johanna Hol Fosse, Grim Rømo). **Corresponding author and email address** Ragnhild Tønnessen.

**Keywords:** highly pathogenic avian influenza, H5N1, cross-species transmission, spillover infection, wildlife-livestock interface, post-outbreak surveillance

## Abstract

A 2023 outbreak of highly pathogenic avian influenza in seabirds in Norway caused substantial environmental contamination of grazing areas frequented by local sheep. Eleven months later, 220 sheep were tested for antibodies to type A influenza and H5 subtype using ELISA, haemagglutination inhibition, and microneutralisation assays. One ewe (0.5%) tested positive by all methods, consistent with prior spillover infection. This underscores the importance of restricting livestock access to outbreak areas to mitigate cross-species transmission and zoonotic risk.

## Manuscript text

### Exposure of grazing sheep to highly pathogenic avian influenza virus (HPAIV) H5N1 during an outbreak in seabirds in Norway, July 2023

Highly pathogenic avian influenza (HPAI) (H5Nx) clade 2.3.4.4b viruses have spread globally since 2020, causing fatal disease in a wide range of wild bird species with increased rates of transmission to mammals (1). In July 2023, more than 15,000 dead seabirds were recorded in association with an outbreak caused by HPAIV H5N1 EA-2022-BB in northern Norway (2). In the early phase of the outbreak the number of carcasses far exceeded the capacity for removal, resulting in substantial contamination of surrounding rough grazing areas frequented by local sheep. Sheep were observed in highly contaminated areas (Figure 1A), including lambs that investigated diseased birds with their muzzle before feeding from their mothers (Figure 1B, Supplementary Video 1). At the time of the outbreak, ruminants were not considered susceptible to HPAI, the sheep appeared healthy, and the potential for spillover infection was not investigated further.

**Figure 1:**
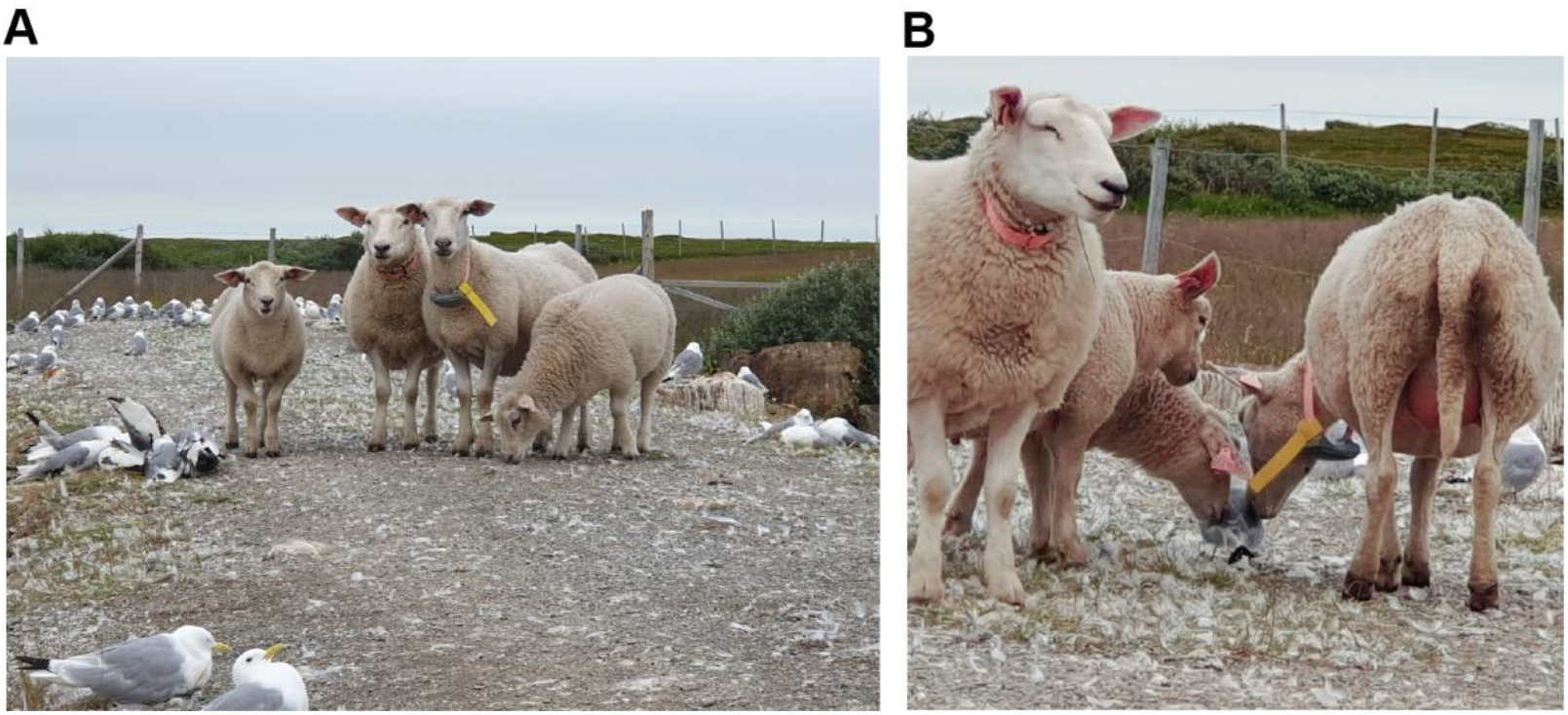
Exposure of sheep during the 2023 HPAI outbreak in seabirds. (A) Sheep surrounded by diseased and dead birds and other contaminated materials. (B) Lamb and ewe in close muzzle contact with diseased bird. Immediately after the photo was taken, the lamb fed from the ewe, illustrating a potential route for direct infection of the udder (Supplementary video 1). Photos: Grim Rømo.

### Detection of antibodies specific to type A influenza and H5 subtype avian influenza in ovine serum collected in June 2024, eleven months after HPAI exposure

In March 2024, an extensive epizootic caused by HPAIV H5N1 was confirmed in dairy cattle in the United States (3, 4), raising the awareness of potential spillovers from wild birds to ruminants. Clinical signs in cows were associated with mastitis, and high virus titres were detected in milk (3-5). Inter-mammalian transmission was documented between cows, from cows to dairy farm workers, and to cats ingesting raw milk (3, 4, 6, 7). In June 2024, blood samples were collected from 220 Norwegian white sheep from the three farms exposed during the 2023 avian outbreak, as part of post-outbreak surveillance. Parallel milk samples were collected from 59 lactating sheep from one of the farms. Serum samples were tested for the presence of antibodies specific to type A influenza nucleoprotein, using a commercial ELISA (ID screen Influenza A Antibody Competition Multi-species ELISA, Innovative Diagnostics, France). The initial analysis, following the manufacturer’s recommendations for bovine serum available at the time (8), detected no positive samples. When samples were re-analysed in May 2025 according to updated recommendations for bovine serum (9), antibodies specific to type A influenza were detected in two ewes, born 2022 and 2023. The sheep born in 2022 (#76) also tested positive for the presence of antibodies specific to haemagglutinin subtype 5 (H5 subtype) avian influenza in a commercial ELISA (ID screen Influenza H5 Competition 3.0 Multispecies ELISA, Innovative Diagnostics, France), following the manufacturer’s recommendations for bovine serum (10). This sample also showed low-to-moderate neutralising activity against a H5 subtype clade 2.3.4.4b virus (A/turkey/Italy/21VIR9520-3/2021 (H5N1)) by the haemagglutination inhibition (HI) test (titres 1:10-1:20) and the microneutralisation (MN) test (titre 1:40), confirming the presence of anti-H5 antibodies. The sample did not inhibit haemagglutination by two different low pathogenic avian influenza (LPAI) H5N3 viruses of Eurasian non-Gs/Gd lineage (A/teal/England/7394/2006 and A/Mallard Duck/Netherlands/38/2008), suggesting that the humoral response was specific for currently circulating 2.3.4.4b clade H5 viruses. The other sheep (#68) that tested positive for type A influenza antibodies gave doubtful results for antibodies to H5 subtype by ELISA and tested negative by both MN and HI. A third (#140) and fourth (#10) sheep tested positive and doubtful for antibodies to H5 subtype by ELISA, respectively. Interestingly, MN titres of 1:20 were recorded in both animals, despite testing negative by both the HI assay and the type A influenza ELISA. Four sheep (#150, 152, 211, 216) tested positive by the MN assay only (titres 1:20), while three other sheep (#3, 65, 120) tested doubtful in the H5 ELISA but negative by all other assays. Milk samples from 59 individuals, including the two that tested positive for antibodies to type A influenza in serum, all tested negative for antibodies to type A influenza and H5 subtype by ELISA and were not analysed further. Figure 2 shows ELISA results of individual sheep, with the HI-positive individual highlighted in red. Data from individuals with positive results in one or more assays are provided in Table 1.

**Table 1.**
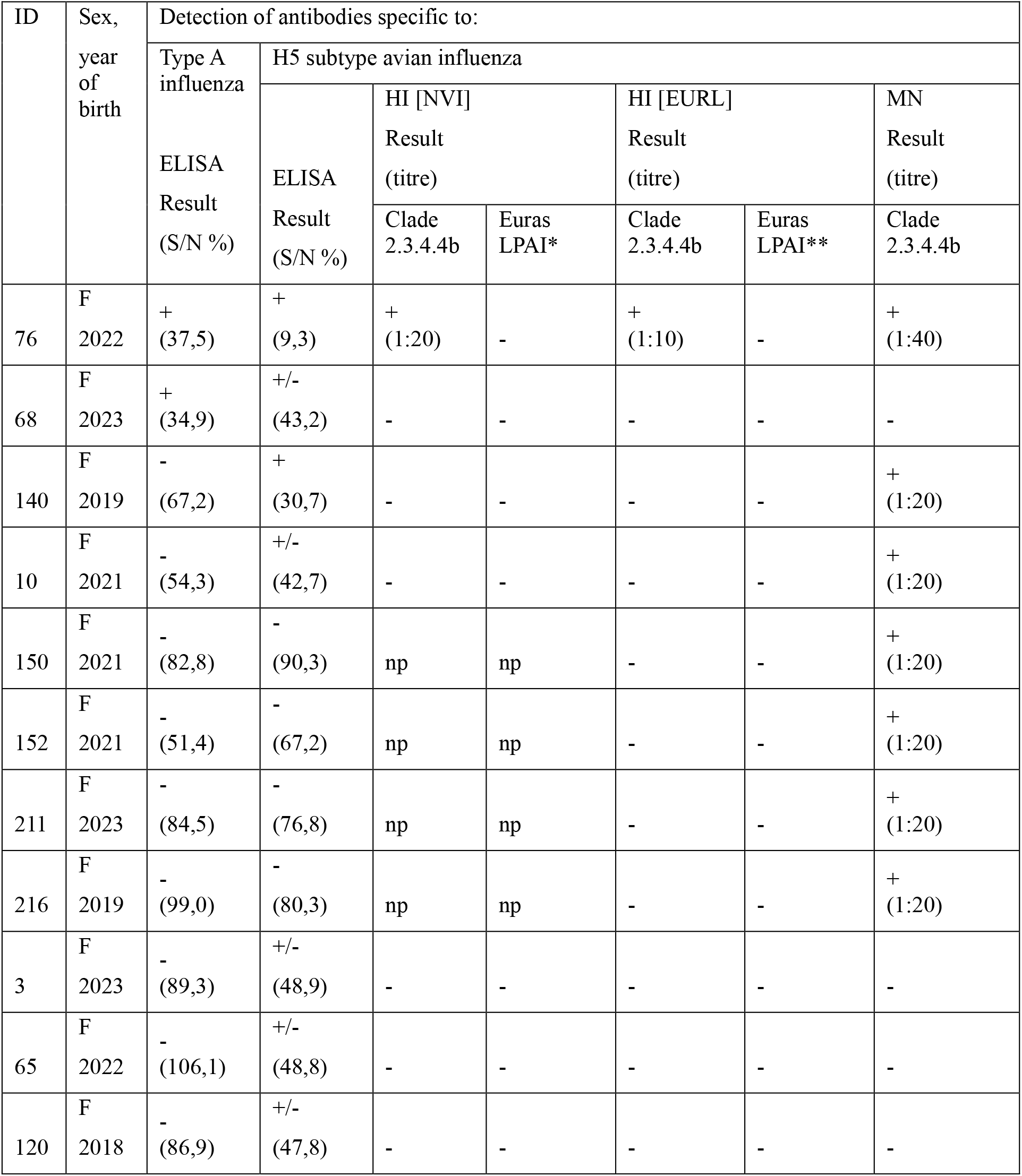
Sheep testing positive in one or more assays for serum antibodies specific to type A influenza nucleoprotein (ELISA = ID screen Influenza A Antibody Competition Multi-species ELISA) or H5 subtype avian influenza (ELISA = ID screen Influenza H5 Antibody Competition 3.0 Multi-species ELISA, HI = haemagglutination inhibition test, MN = microneutralisation assay, Clade 2.3.4.4b = A/turkey/Italy/21VIR9520-3/2021 (H5N1), Euras LPAI (Eurasian low pathogenic avian influenza) = *A/teal/England/7394/2006 (H5N3) or **A/Mallard Duck/Netherlands/38/2008 (H5N3), NVI = Norwegtian Veterinary Institute, EURL = European Reference Laboratory for Avian Influenza and Newcastle disease, Istituto Zooprofilattico Sperimentale delle Venezie, np = not performed).

**Figure 2:**
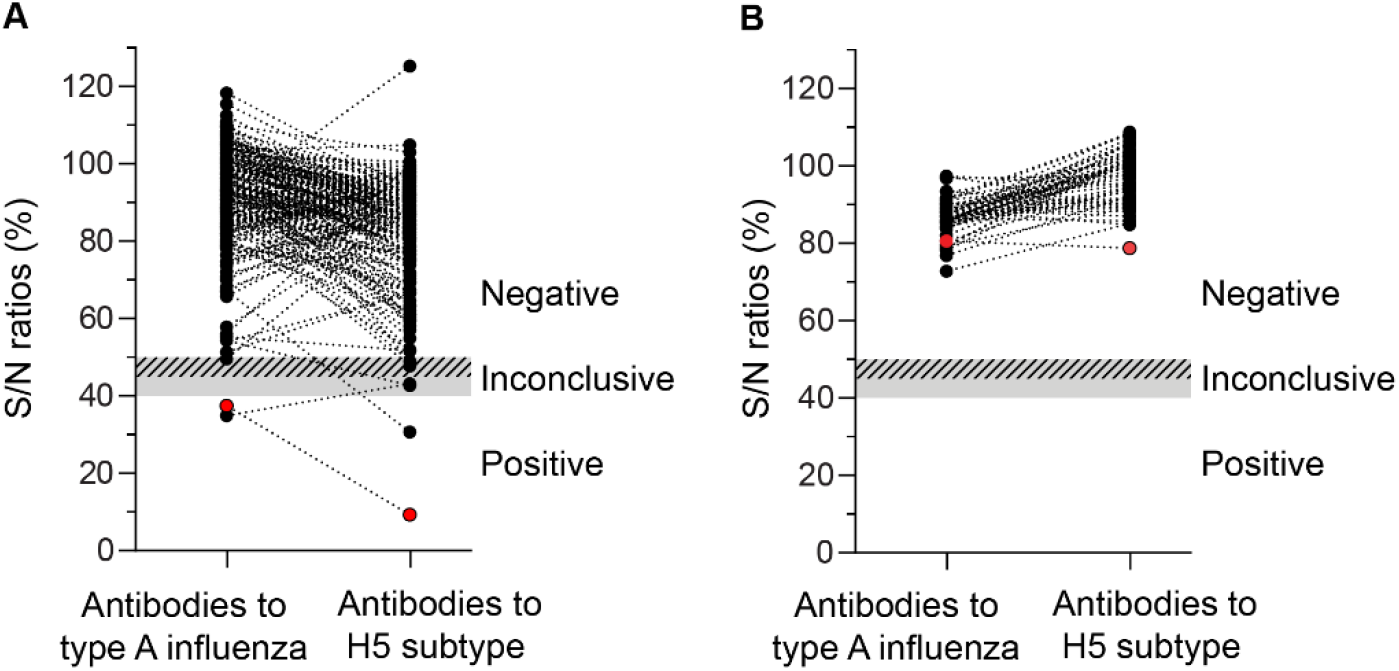
Detection of antibodies to type A influenza and H5 subtype avian influenza in sheep following HPAI exposure. (A) Serum (n=220) and (B) milk (n=59) samples were analysed for the presence of antibodies specific to type A influenza nucleoprotein and H5 subtype avian influenza using commercial ELISA kits (ID screen Influenza A Antibody Competition Multi-species ELISA and ID screen Influenza H5 Antibody Competition 3.0 Multi-species ELISA, both from Innovative Diagnostics, France), in accordance with the manufacturer’s recommendations for bovine samples (9, 10). Dots show individual S/N-ratios with lines connecting results from the same sheep. The inconclusive ranges are indicated as hatched (type A influenza) and grey (H5 subtype) areas. The sheep with positive results in HI and MN serum assays is highlighted in red.

## Discussion

We here document the detection of antibodies specific to type A influenza and H5 subtype avian influenza across multiple assays in a single sheep in Norway. This individual was one of 220 sheep with documented exposure to substantial environmental contamination during a major HPAI outbreak in seabirds eleven months earlier, in 2023. Considering the degree of exposure to contaminated material during the outbreak, the young age of the sheep, and the limited exposure to free-range grazing in the period between the outbreak and sampling, it is likely that the antibody response originated from this specific exposure event.

Detection of antibodies in one out of 220 sheep corresponds to a serological prevalence of approximately 0.5%. It is possible that this number underestimates the true rate of spillover infections, as the eleven-month interval between exposure and sampling could have decreased the sensitivity of our surveillance. Six additional sheep showed partial serological evidence of H5 exposure, based on detection of antibodies against the H5 virus in the ELISA or the MN assay, despite testing negative in the comparatively less sensitive HI assay and for antibodies against type A influenza. These findings should be interpreted with caution, considering the gaps in knowledge of antibody response kinetics to different viral proteins following H5 avian influenza virus infection in sheep and the limited validation of serological assays in this species. For comparison, cows infected with HPAIV H5N1 mount antibody responses to influenza A nucleoprotein and H5 in both milk and serum within 1-2 weeks of infection (11), but the duration of these responses remains undetermined. No antibodies were detected in milk in our study, suggesting that serum may be a more sensitive matrix than milk for serological HPAI surveillance in sheep.

Active HPAIV H5N1 infection was recently documented by Banyard *et al.* in a sheep in Great Britain, on a farm with an HPAI outbreak in captive birds (12). Despite the absence of clinical signs, viral RNA was detected in milk, suggesting that the tropism of HPAIV in sheep may resemble that observed in cattle (3, 11). In this sheep, HI titres peaked at 1:160, considerably higher than the titre detected in our investigation. Antibody titres are strongly influenced by the time that has passed since the last exposure. Serological investigations in two human patients with confirmed H5N1 infection showed that MN and/or HI titres in serum samples collected 5 to 18 months post-infection were reduced by eight-fold or more compared to peak levels (13, 14). If a similar waning pattern applies to sheep, the HI titres detected in our investigation would be consistent with a peak response comparable to that observed by Banyard *et al.* (12). Moreover, in their discussion of potential routes of infection, Banyard *et al.* suggest that infective material could have been introduced into the teat by suckling lambs. Our observations of lambs exploring diseased birds with their muzzles before feeding from their mothers support the relevance of this hypothesis, although no conclusions about entry point or infection site can be made from our investigation.

As our study is based on retrospective serological investigations, we cannot exclude the possibility that the observed antibody response was triggered by prolonged mucosal exposure to high antigen loads in the absence of active infection. However, in the absence of viral replication, the induction of a humoral response typically requires mechanisms that allow the antigen to cross the epithelium, such as parenteral administration or the use of adjuvants (15, 16). Therefore, we propose that actual spillover infection remains the most plausible explanation for the detection of antibodies against both surface and internal viral proteins in a sheep with documented exposure to diseased birds in a highly contaminated environment. Our findings suggest that HPAI spillovers from wild birds to sheep can occur given sufficient exposure, although the frequency appears to be low, even under conditions of high infection pressure.

HPAI in ruminants constitutes a zoonotic threat, with cattle identified as the likely source of 41 human cases during the ongoing U.S. outbreak (7). Our findings add to previous reports providing direct and indirect evidence of HPAI spillover from wild birds to small ruminants (12, 17). From a One Health perspective, our findings underscore the need to prevent livestock contact with HPAI-contaminated grassland, diseased birds, and carcasses, and to include small ruminants in HPAI outbreak investigation and surveillance.

---

**Supplementary video 1:** Complementary to Figure 1. Footage showing lamb feeding from ewe immediately after close muzzle contact with a diseased seabird during the 2023 HPAI outbreak in northern Norway. Video by Grim Rømo.

## Author statements

### Author contributions

**JHF, GR**, and **RT** conceptualised the study, interpreted results, and jointly developed the final manuscript. **GR** initiated contact with farms and local authorities and organised and carried out sample and epidemiological data collection. **JHF** planned and coordinated the serological analyses, analysed and visualised data, and wrote the first draft of the manuscript. **RT** managed the overall project. **FB** provided expertise and conducted microneutralisation tests and confirmatory analyses. **KS** contributed to sample collection and processing. **IKM** and **KU** performed the serological analyses. All authors reviewed the drafts and approved the final manuscript.

### Ethical approval

Blood sampling was performed as part of post-outbreak surveillance and falls within the definition “non-experimental clinical veterinary practices”, which is not considered scientific research and hence does not require prior approval by the national animal research authority (DIRECTIVE 2010/63/EU).

### Consent for publication

All participating farmers provided written informed consent for inclusion of their animals and related information in this publication.

### Data statement

The full set of raw data from the study have been deposited in the diagnostic record system of the Norwegian Veterinary Institute, where they are stored in accordance with institutional and national data management protocols. Anonymised data can be provided upon request.

## Acknowledgements

We thank the farmers for their invaluable cooperation in providing information and for assistance with sample collection. We are also grateful to Gørill Hogseth (Norwegian Food Safety Authority, Norway) for her assistance with sample collection, and to Francesca Ellero (Istituto Zooprofilattico Sperimentale delle Venezie, Legnaro, Italy) for her support in coordinating sample shipment logistics.

